# Macroscale brain states support the control of semantic cognition

**DOI:** 10.1101/2024.02.29.582250

**Authors:** Xiuyi Wang, Katya Krieger-Redwood, Yanni Cui, Jonathan Smallwood, Yi Du, Elizabeth Jefferies

## Abstract

Understanding how the human brain adapts to varying cognitive demands is crucial in neuroscience. Here, we examined how networks involved in controlled semantic retrieval reconfigure themselves to generate neurocognitive states appropriate to different task contexts. We parametrically varied the demands of two semantic tasks - global association and feature matching judgments - and contrasted these effects of cognitive control with those of non-semantic tasks. We then characterized these effects on the cortical surface and within a whole-brain state space, anchored by the top three dimensions of intrinsic connectivity. Our results revealed that demanding semantic association tasks elicited more activation in the anterior regions of the prefrontal and temporal cortex. In contrast, difficult semantic feature matching tasks produced more posterior activation, aligning closely with regions engaged during multiple demanding non-semantic tasks. In both semantic feature matching and non-semantic contexts, the difficulty effects were situated towards the controlled end of a dimension capturing functional separation between cognitive control and default mode regions. Conversely, in semantic association tasks, the difficulty effects elicited similar responses across both cognitive control and default mode networks. Furthermore, controlled association and non-semantic control were located towards the heteromodal end of a heteromodal- unimodal dimension, while semantic feature matching involved a brain state that was more visual and unimodal. These findings demonstrate that a variety of brain states underpin controlled cognition. Specifically, cognitive control regions *interact* with heteromodal semantic knowledge system to identify contextually relevant conceptual overlaps (e.g., associating ’DOG’ with ’BEACH’), and *separate* from heteromodal memory regions for modality-specific conceptual overlaps (e.g., connecting ’DALMATIAN’ with ’BLACK AND WHITE’).

## Introduction

Adaptive behavior hinges on understanding the meanings of our surroundings and modulating our responses accordingly. While research has focused on how the brain stores semantic information and controls cognition to achieve our goals, fewer studies have investigated the intersection of these domains to understand how we flexibly retrieve context-appropriate information. For example, searching for your dog on a crowded beach might focus on visual features like color and shape. In contrast, at a family gathering, associative details become more relevant – recognizing that dogs are strongly food-motivated, and chocolate is harmful to them. These scenarios highlight our ability to adapt semantic retrieval to different situations. However, current descriptions of brain networks underpinning conceptual representation and control fall short in explaining how we generate diverse brain states that can support these different retrieval patterns.

Semantic cognition relies on conceptual representations distilled from sensory-motor features within heteromodal hub(s), including anterior temporal cortex, as well as two networks that support cognitive control – the semantic control network (SCN) and multiple demand network (MDN) (Lambon Ralph et al., 2017; Xu et al., 2016). The MDN, particularly its frontoparietal regions including the bilateral inferior frontal sulcus and intraparietal cortex, responds to executive demands across various tasks (Assem et al., 2022, 2020; Duncan, 2010; Fedorenko et al., 2013). It is thought to support domain-general control processes, such as maintaining goals applicable to different types of representations, including semantic information (Duncan, 2010). Concurrently, meta-analyses of semantic tasks reveal a partially-overlapping yet dissociable set of SCN regions, including the left inferior frontal gyrus (IFG), posterior temporal cortex (PTC), and dorsomedial prefrontal cortex (dmPFC) (Jackson, 2021; Noonan et al., 2013). These regions show stronger activation when there is an increased necessity to constrain conceptual retrieval, for example, to access weaker associations, ambiguous relationships or specific features not strongly linked to a concept (Jackson, 2021; Noonan et al., 2013). The SCN is engaged in controlled, flexible semantic retrieval but is less activated by demanding non-semantic tasks (Chiou et al., 2023; Gao et al., 2021; Gonzalez Alam et al., 2018; Wang et al., 2020). Semantic and non-semantic controlled states also differ in lateralization: the SCN is primarily left-lateralized, whereas the MDN is bilateral (Fedorenko et al., 2013; Jackson, 2021; Noonan et al., 2013).

Given that MDN is recruited across domains and SCN is implicated in diverse semantic tasks, a pivotal question emerges: how do we generate whole-brain states to focus on different aspects of knowledge fitting a specific task context (Greene et al., 2023)? A clue lies in the relationship between these two control networks. Although proximal on the cortical surface, for example, in the left lateral prefrontal cortex, they occupy distinct positions in a hierarchy from sensory-motor to heteromodal cortex (Chiou et al., 2023; Wang et al., 2020). This proximity might elucidate why controlled semantic retrieval elicits stronger responses in the left anterior lateral prefrontal cortex, while non-semantic control effects and semantic feature matching activate the posterior lateral prefrontal cortex (Badre et al., 2005; Badre and Wagner, 2007; Gold et al., 2006; Pang et al., 2023). These functional differences might reflect the principal dimension of intrinsic connectivity, which explains the largest variance in resting-state fMRI and differentiates between heteromodal and unimodal processing. Prior research suggests that SCN is closer to the heteromodal end of this dimension than MDN (Wang et al., 2020). This leads to the prediction that difficulty effects in semantic association and semantic feature matching will not only show topographical differences in the left lateral prefrontal cortex but that these differences will extend to anterior and posterior areas of posterior temporal and medial prefrontal areas, where SCN and MDN are adjacent (Jackson, 2021; Noonan et al., 2013).

Different states of controlled cognition may reflect specific configurations of large-scale brain networks, which can be characterized in terms of multiple dimensions of intrinsic connectivity (Bolt et al., 2022; Margulies et al., 2016). In addition to the principal dimension of intrinsic connectivity differentiating heteromodal from unimodal processing, a second dimension separates visual from auditory-motor processes, while a third dimension delineates the functional separation between the Default Mode Network (DMN) and cognitive control systems (Bolt et al., 2022; Margulies et al., 2016). When controlled semantic retrieval is required to establish relevant thoughts and behaviors in the absence of an externally-imposed goal (for example, when we focus on weak associations relevant to the context), heteromodal regions that support long-term semantic knowledge are thought to be integrated with control processes that can shape retrieval to suit the circumstances (Davey et al., 2016; Luppi et al., 2024, 2022; Wang et al., 2020). Conversely, controlled non-semantic states are associated with anti-correlation between control and DMN networks. By mapping controlled activation patterns within a whole-brain state space defined in terms of the first three dimensions of variation in intrinsic connectivity, spatial activation differences across the whole brain can be explained in terms of their reliance on heteromodal versus unimodal cortex, visual versus auditory-motor inputs, and the extent to which control networks are engaged without DMN. Consequently, this approach allows us to understand diverse patterns of network interactions across different task contexts.

In this study, we explored how networks implicated in control are engaged on the cortical surface and in a whole-brain state space defined by the top three dimensions of intrinsic connectivity. To achieved this, we parametrically varied the demands of two semantic tasks—global association and semantic feature matching—and contrasted the effects of control with those of two non-semantic tasks. Specifically, in the association task, participants retrieved global associations using a broad range of semantic features. Conversely, in the semantic feature matching task, they made decisions about words based on visual attributes like color, specified by a task instruction that provided an explicit goal. Task difficulty was manipulated by altering the strength of associations and feature similarity for word pairs, respectively. We then compared activation patterns for these semantic control aspects with those in more challenging spatial working memory and math judgments. Our study had three primary objectives: (i) To establish if brain regions supporting controlled retrieval of semantic associations are anterior to those for visual feature selection (cf. Badre et al. 2005), but extending beyond the left inferior frontal gyrus to include medial prefrontal and posterior temporal cortex, thereby indicating an organized topographical dissociation in whole-brain organization. (ii) To determine whether control processes linked to semantic feature matching overlap more with non-semantic control regions than those engaged in the controlled retrieval of semantic associations. (iii) To understand the organization of cognitive control in neural state space, in which differences in activation are interpreted in terms of dimensions of whole-brain functional organization. Thus, our research builds on prior findings of multiple control networks (SCN versus MDN) and functional dissociations within LIFG, to establish whether multiple modes of controlled cognition are underpinned by distinct dimensions of neural organization.

## 2. Results

This study analyzed two datasets collected at the University of York, UK. The first dataset involved two semantic control tasks (Wang et al., 2023), while the second dataset involved two non-semantic control tasks, aimed at localizing the MDN (Wang et al., 2021, 2020).

### 2.1. Behavioral data

#### 2.1.1. Semantic tasks

We parametrically manipulated the difficulty of two semantic tasks (Fig. 1). In these tasks, participants decided whether a word pair shared a semantic relationship by making Yes/No decisions based on either: (i) association strength, accessing whether two concepts were globally related in meaning; or (ii) feature overlap, evaluating whether two concepts shared similar visual features (either color or shape). The semantic association task presented word pairs with varying degrees of association. Stronger associations were expected to facilitate decision making for related (“Yes”) trials, since they are typically more easily accessible from the semantic long-term store. Conversely, relatively strong associations could complicate unrelated (“No”) decisions (see supplementary section 1.2). In this task, participants were not given an explicit goal or specific instructions on how to link the concepts but were asked to make decisions based on overall semantic similarity. This design directed controlled retrieval towards aspects of the concepts that matched a shared context, with information from the semantic store providing this context.

**Fig. 1.**
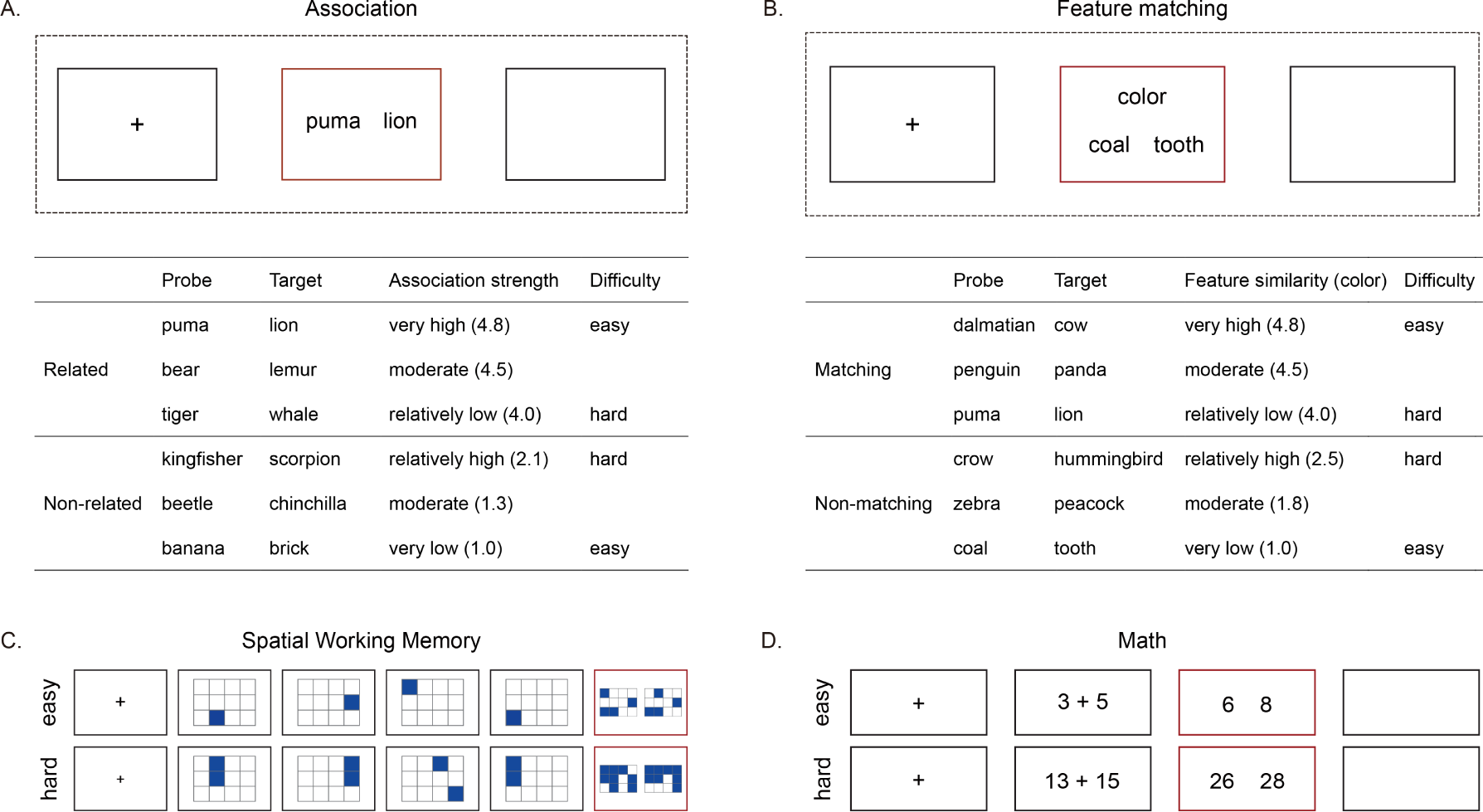
Illustration of the semantic and non-semantic tasks. A – Semantic association task: Participants made yes/no decisions about whether pairs of words were globally semantically associated or not. We parametrically manipulated the association strength between the probe and target word, typically judged to be related or unrelated on a 5- point rating scale. B – Semantic feature matching task: Participants decided if probe and target concepts shared a specific visual semantic feature (color or shape), indicated at the top of the screen during each trial. The feature prompt, probe and target words appeared simultaneously. We parametrically manipulated the degree of feature similarity between the probe and target concepts that were typically judged to be matching or non-matching for the specified feature on a 5-point rating scale. C and D – Non-semantic tasks for domain-general control: C involved a spatial working memory task where participants tracked sequentially presented locations. D entailed math decision tasks, requiring the maintenance and manipulation of single or double-digit numbers.

In the semantic feature matching task, in contrast, participants were asked to decide if two concept words shared a specific visual feature – color or shape. The word pairs parametrically varied in feature similarity – i.e., how similar the concepts were in terms of the feature being matched. A high degree of feature similarity was anticipated to ease the decision-making for matching (“Yes”) trials, as it would likely increase participants’ confidence in their matching decisions. Conversely, lower feature similarity was expected to simplify non-matching (“No”) trials, making the basis for non-matching decisions more apparent (see supplementary section 1.3). Unlike the semantic association task, the semantic feature matching task explicitly required participants to focus on and execute a specific semantic goal for semantic retrieval, making broader conceptual information about the concepts irrelevant.

Our first analysis verified the effectiveness of our parametric manipulation of task demands. To examine how semantic association strength influenced response time (RT) in the semantic association task, we built a linear mixed effect model. This model accounted for individual differences in the difficulty effect by including random intercepts and slopes. We compared a model incorporating a linear effect of semantic association strength with a model without this effect. The results showed that association strength significantly facilitated decision making for related trials (z = -9.244, p < 0.0001) but had no discernible effect on unrelated trials (z = 0.018, p = 0.986), after controlling for feature similarity and global similarity, the latter being the overall similarity of each word pair as rated by an independent group of 30 participants (Fig. 2A). We conducted a comparable analysis for the feature matching task to investigate how feature similarity influenced response times and accuracy. The results indicated that higher feature similarity facilitated decision-making for matching trials (RT: z = -10.51, p < 0.0001), but impeded decisions for non-matching trials (RT: z = 11.97, p < 0.0001) after controlling for association strength and global similarity (Fig. 2B and 2C).

**Fig. 2.**
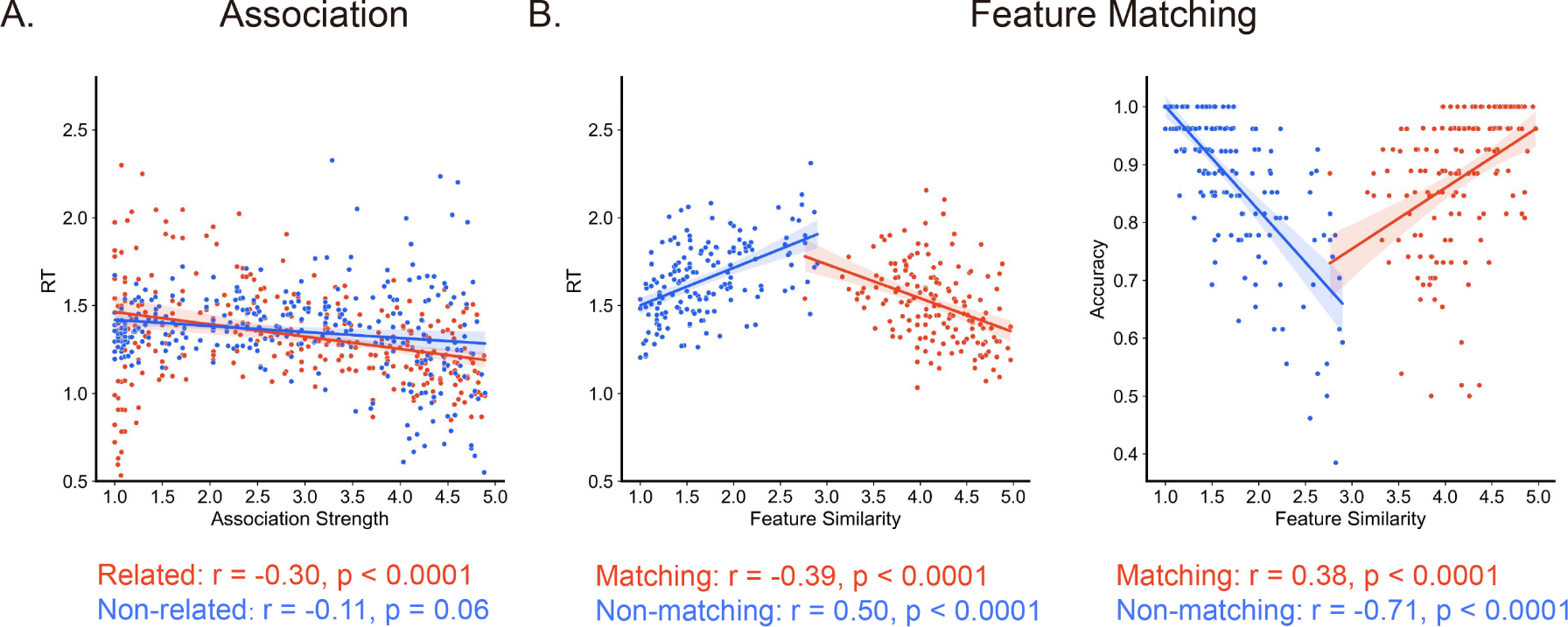
Behavior data for the semantic tasks. A – In the semantic association task, semantic association strength was negatively correlated with response time for the related trials, but had no significant correlation for the unrelated trials. B – In the feature matching task, feature similarity was negatively correlated with RT for the matching trials, but positively correlated for the non-matching trials. C – In the feature matching task, feature similarity showed a positive correlation with accuracy for matching trials, but a negative correlation for the non-matching trials. An analysis of accuracy for the association matching task was not performed because participants made their own judgements about which words were related and which were unrelated. For trials with intermediate association strengths, these decisions vary across individuals.

#### 2.1.2. Non-semantic tasks

To investigate the overlap between effects of semantic control in the two semantic tasks and domain-general cognitive control, we included two non-semantic tasks commonly used to localize regions of the MDN: a spatial working memory task and a math task (Fedorenko et al., 2013). In the spatial working memory task, participants tracked locations presented in sequence, with the easy version involving one location per slide and the hard version two locations, thus increasing working memory load. In the more demanding version, both accuracy and RT were affected, showing decreased accuracy (t (26) = -8.97, p = 7.31 * e-10) and increased RT (t (26) = 7.14, p = 7.20 * e-8) compared to easier trials. Similarly, the math task ranged from single-digit additions in the easy version to double-digit additions in the hard version. The more demanding condition resulted in lower accuracy (t (26) = -6.73, p = 2.19 * e-7) and longer RTs (t (26) = 12.06, p = 8.04 * e-13) compared to easier trials. These contrasts between hard and easy versions of the tasks have been utilized to identify MDN regions responsive to cognitive control demands (Fedorenko et al., 2013; Wang et al., 2021, 2020).

### 2.2. Effects of strength of association and feature similarity on brain responses

Next, we evaluated whether our difficulty manipulations in the semantic association and feature matching tasks engaged common or distinct brain regions. First, we investigated whether the spatial differences in the left IFG previously reported — more anterior activation for global association matching and more posterior for feature matching (Badre et al., 2005) — would be replicated with our parametric difficulty manipulation in these two tasks. Secondly, we explored whether this functional dissociation extended to other brain areas, such as the left posterior temporal and medial prefrontal regions. Confirmation of this would indicate that adjacent yet functionally distinct large-scale neural networks are systematically organized on the brain’s surface, with each supporting different facets of semantic control.

We pinpointed brain regions that exhibited a stronger response to more difficult trials in the two semantic tasks. This increase in activation occurred when (i) association strength was lower for related ’Yes’ trials or higher for unrelated ’No’ trials in the semantic association task, and (ii) feature similarity was lower for matching ’Yes’ trials or higher for non-matching ’No’ trials in the feature matching task. We also identified regions that showed greater activation in easier trials. The main task effects (i.e., greater activation during the task relative to the resting baseline) are shown in the Supplementary Materials (Fig. S1).

Fig. 3A shows the parametric manipulation of semantic association strength (p < 0.05, FDR-corrected), and Fig. 5E shows the corresponding unthresholded map. Multiple regions showed positive effects of decision difficulty, with increased BOLD response when association judgements were more difficult, including temporal-occipital cortex, intraparietal sulcus, inferior frontal sulcus and pre-supplementary motor area (Fig. 3A). Negative effects of this variable, reflecting a stronger BOLD response during easier association judgments, were found in default mode network regions in lateral anterior-to-mid temporal cortex, angular gyrus, and medial and superior frontal regions (Fig. 3A). The unthresholded maps for difficulty effects in related and unrelated trials were spatially similar (Fig. S2, i.e., the effects of weaker associations when items were judged to be related and stronger associations when items were judged to be unrelated were significantly correlated using spin permutation).

**Fig. 3.**
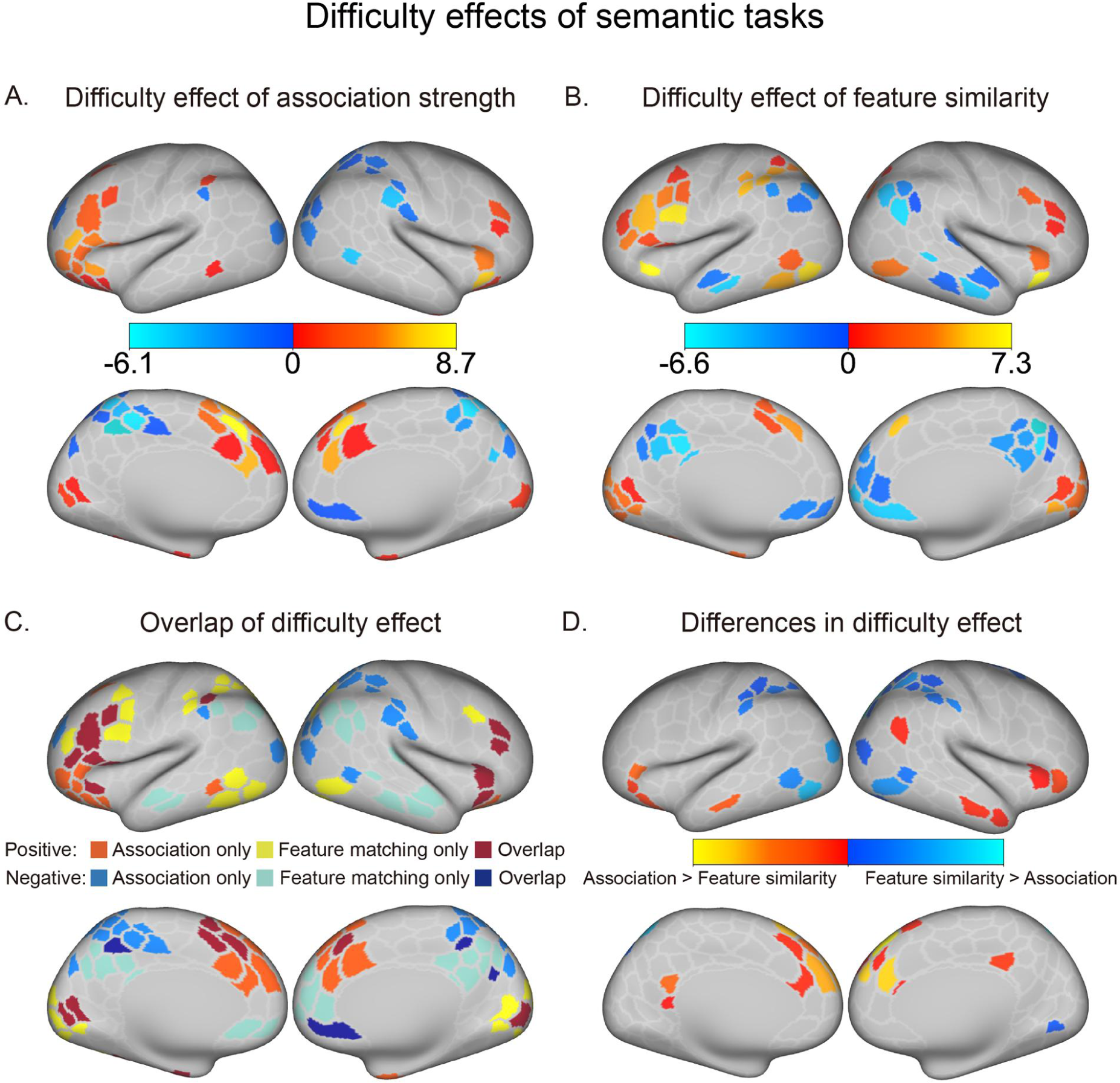
The parametric difficulty effects of semantic association and feature similarity, and their comparison. A – The effect of decision difficulty in the semantic association task. Warm colors indicate regions with increased activation during more difficult trials (i.e., weaker association strength in associated trials and stronger in non-associated trials). Cold colors represent the regions that showed the reverse trend (i.e., showing greater activation in less demanding trials). B – The effect of decision difficulty in the semantic feature matching task. Warm colors mark regions with heightened activation for more difficult trials (i.e., lower feature similarity in matching trials and higher in non-matching trials). Cold colors denote regions showing the opposite trend. C – Overlap in decision difficulty effects for these two tasks. For semantic association, increased difficulty elicited stronger activation in anterior cortex, while in feature similarity, it led to stronger engagement in posterior cortex. D – The comparison of the difficulty effects in these two tasks. Warm colors denote regions more strongly modulated by association strength compared to feature similarity, and cold colors indicate areas showing the opposite pattern.

Fig. 3B shows the thresholded difficulty effect of feature similarity (p < 0.05, FDR-corrected) and Fig. 5F shows the corresponding unthresholded map. Positive effects of decision difficulty across matching and non-matching trials (i.e., stronger responses to harder trials) were found in inferior frontal sulcus, pre-supplementary motor area, temporal-occipital cortex, and intraparietal sulcus (Fig. 3B). Conversely, regions in the DMN showed negative effects of decision difficulty (i.e., stronger responses to easier trials), including lateral anterior-to-mid temporal cortex, angular gyrus, medial and superior frontal regions, and posterior cingulate cortex (Fig. 3B). The unthresholded maps for difficulty effects in matching and non-matching trials were spatially similar (Fig. S2; i.e., the effects of lower similarity for matching trials and higher similarity for non-matching trials were correlated using spin permutation).

Although there was considerable overlap in the effect of difficulty for association strength and feature similarity (Fig. 3C), there were also differences in difficulty effects across tasks (Fig. 3D). A direct comparison of the parametric difficulty effects in semantic association and feature matching tasks revealed stronger modulation by difficulty in the semantic association task within DMN regions, including the posterior cingulate cortex, ventral prefrontal cortex, and temporal pole (Fig. 3D). This aligns with the view that the semantic association task more intensively engages controlled retrieval from heteromodal regions. Conversely, stronger modulation by difficulty in the semantic feature matching task was found in cognitive control regions, such as the intraparietal sulcus (IPS), inferior parietal lobule (IPL), and temporal-occipital cortex showed (Fig. 3D). We found that responses to difficulty in global association were more anterior compared to feature matching in the left lateral prefrontal, medial prefrontal, and left posterior temporal cortex, (Fig. 3C). This finding demonstrates that task difficulty can be differentiated not only by activation within individual regions but also by whole-brain topography. The increased demand in feature matching trials might rely more on the controlled retrieval of sensory information to focus on specific visual features of a concept, thus eliciting stronger activation in the lateral and polar occipital cortex. Conversely, more difficult semantic association tasks may predominantly depend on the controlled retrieval of heteromodal long-term knowledge, as they require establishing a linking context for the two words based on general semantic information. This could explain the more anterior response in regions more physically further from the sensory-motor cortex.

### 2.3. Comparison of semantic and non-semantic task demands

To assess the overlap between difficulty effects in semantic tasks and brain regions responsive to non-semantic task demands, we conducted three analyses. First, we compared hard with easy versions of spatial working memory and math judgements (thresholded maps in Fig. 4A and 4B, unthresholded maps in Fig. 6A and 6B). Fig. 4C and 4D illustrate the extent of overlap between the difficulty effects of semantic tasks and non-semantic tasks. Specifically, 32% of brain regions in the semantic association task overlapped with non-semantic control regions that showed hard versus easy activation in either spatial working memory or math tasks (purple in Fig. 4C), while 71% of parcels in the semantic feature matching task showed this pattern of overlap (blue in Fig. 4D). Next, we defined MDN regions by pinpointing areas that showed a difficulty effect in both spatial working memory task and math task (Fig. 4E). We then compared the activation associated with task difficulty in these MDN regions for the semantic association and semantic feature matching tasks. The difficulty effect was more pronounced for feature similarity than for association strength (t (27) = 7.28, p = 9.91 × 10 e -8; Fig. 4E). Finally, we computed spatial correlations between unthresholded difficulty effect maps for non-semantic tasks (Fig. 6A and 6B) and semantic tasks (Fig. 5E and 5F) and compared these correlations. Non-semantic difficulty showed stronger positive correlation with task demands in feature matching (spatial working memory: left hemisphere (LH): r = 0.72, right hemisphere (RH): r = 0.60; math task: LH: r = 0.68, RH: r = 0.62; all p values = 0) than in semantic association (spatial working memory: LH: r = 0.31, RH: r = 0.08; math: LH: r = 0.21, RH: r = 0.05), with significant differences between these correlations (differences with spatial working memory: LH: z = 5.83, RH: z = 6.08; differences with math: LH: z = 6.11, RH: z = 6.70; all p values = 0). All p-values were FDR-corrected following spin permutation. These findings confirm that the difficulty effect in the feature matching task overlapped more with neural processes implicated in non-semantic control than the semantic association task.

**Fig. 4.**
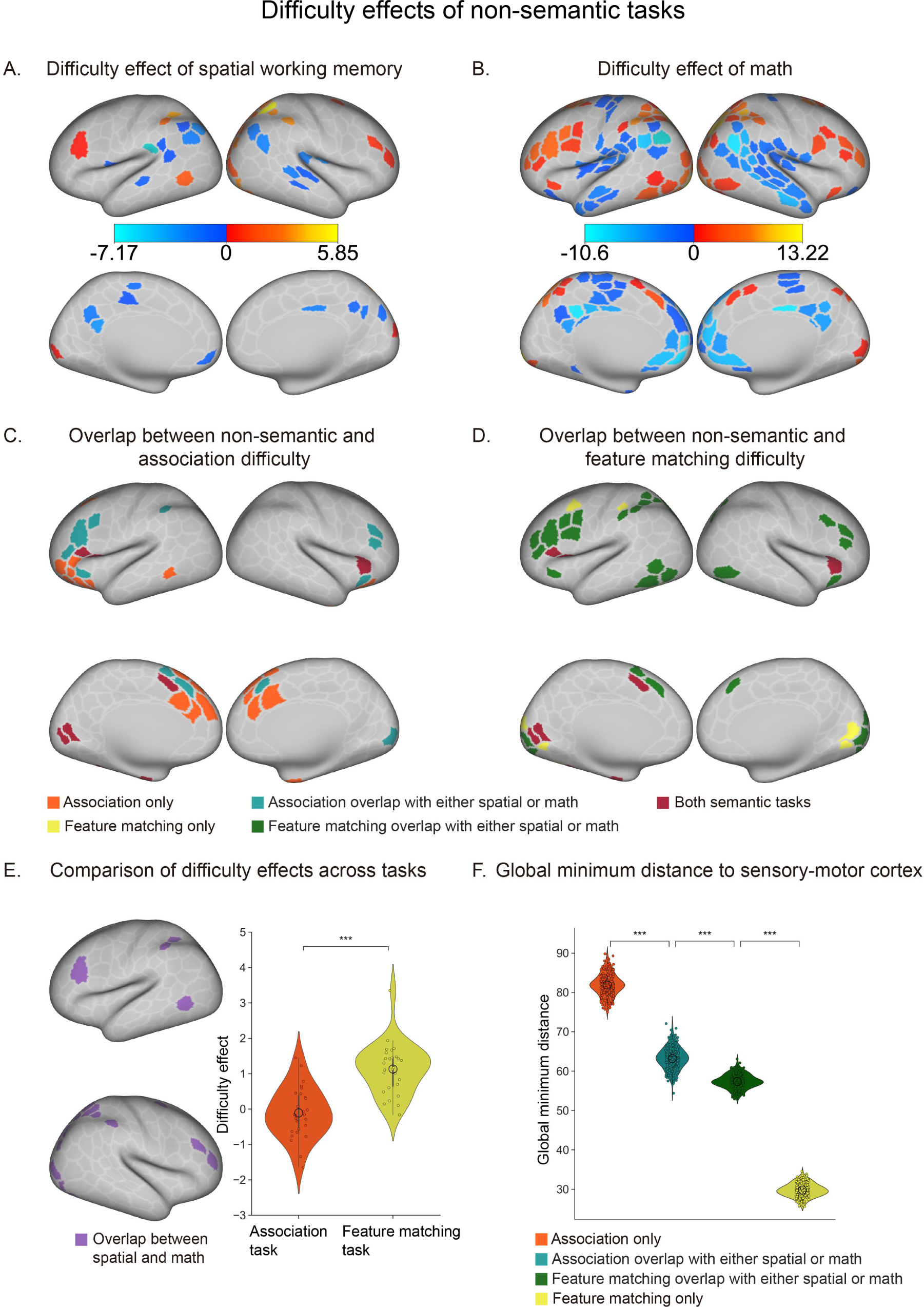
Difficulty effects of spatial working memory and math tasks and their intersection with semantic tasks. A and B – Difficulty effects in spatial working memory and math tasks, respectively. Warm colors indicate regions with increased activation during harder trials (p < 0.05, FDR-corrected), while cold colors show regions with greater activation in easier trials. C – Overlap of regions with positive difficulty effects in the semantic association task (orange) and those responsive to non-semantic control demands (turquois). D – Overlap of regions with positive difficulty effects in the semantic feature matching task (yellow) and those responsive to non-semantic control demands (green). Red regions indicate difficulty effects present in both semantic tasks but not in the non-semantic tasks. E – Greater difficulty effect in semantic feature matching compared to semantic association task within MDN regions (i.e., overlapping regions showing positive effects of difficulty in both spatial working memory and math tasks). F – The global minimum distance to sensory-motor cortex for four types of parcels in C and D, each exhibiting a different pattern of difficulty across tasks. These groups of parcels showed a gradient in their distance from sensory-motor cortex: association-only parcels were the most distant, followed by association and non-semantic parcels, then feature and non-semantic parcels, with feature-only parcels being the closest.

We further examined if the difficulty of semantic association difficulty elicits more anterior brain responses within parcels more physically distant from sensory-motor cortex than semantic feature matching. We analyzed the proximity of these responses to the sensory-motor cortex (Fig. 3). We categorized parcels into four distinct groups based on their response to difficulty: (i) parcels responsive to difficulty solely during the semantic association task (orange in Fig. 4C), (ii) parcels showing difficulty effects in both semantic association and non-semantic tasks (purple in Fig. 4C), (iii) parcels showing difficulty effects in both feature matching and non-semantic tasks (blue in Fig. 4D), and (iv) parcels responsive only to difficulty during the semantic feature matching task (yellow in Fig. 4D). We then computed the global minimum distance from each parcel to its nearest sensory-motor landmarks for each participant (see Method 4.6 for detailed information). These four groups of parcels exhibited a decreasing distance from sensory-motor cortex: association-only parcels were furthest away, followed by association and non-semantic parcels, then feature and non-semantic parcels, and finally, feature-only parcels were the closest to sensory-motor cortex (association-only versus association and non-semantic: t (244) = 118.32, p = 1.53 * e -217; association and non-semantic versus feature and non-semantic: t (244) = 51.94, p = 6.48 * e-134; feature and non-semantic versus feature-only: t (244) = 210.68, p = 5.18 * e -278). All p-values are FDR-corrected. These findings show that the difficulty of semantic associations prompts a more anterior response in regions further from the sensory-motor cortex compared to feature matching.

### 2.4. Situating semantic control effects in a brain state space defined by the dimensions of intrinsic connectivity

The analyses above show that the difficulty effects in semantic association and feature matching tasks exhibit distinct topographical patterns. To reveal how these diverse control processes are organized on the cortical surface, we examined how neural patterns related to task difficulty were situated in a whole-brain state space. This space was defined by the top three dimensions of intrinsic connectivity, identified from resting-state functional MRI data of 245 participants in the S900 release of the HCP dataset, who completed four resting-state scans. Consistent with prior research (Mckeown et al., 2020; Shao et al., 2022; Wang et al., 2020), we focused on the first three connectivity dimensions, which showed the largest eigenvalues (as seen in Fig. 5D scree plot). The first dimension, explaining the most variance (12.75%), separated unimodal (purple-blue in Fig. 5A) from transmodal regions (red-white in Fig. 5A). The second dimension, accounting for 11.29% of the variance, separated somatomotor from auditory cortex (purple-blue in Fig. 5B) from visual cortex (red-white in Fig. 5B). The third dimension, explaining 3.98% of the variance, separated FPCN regions (purple-blue in Fig. 5C) from DMN regions (red-white in Fig. 5C).

**Fig. 5.**
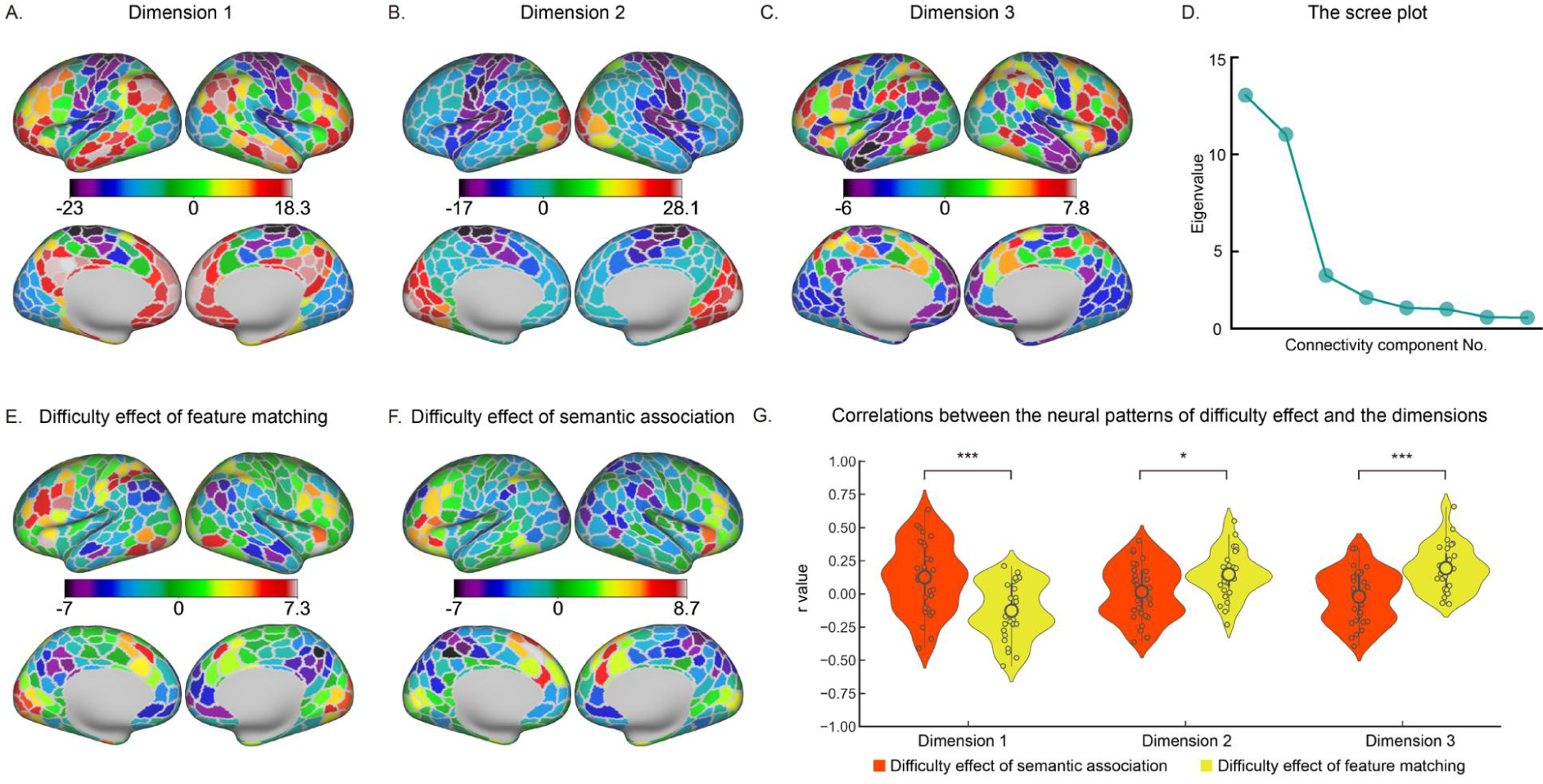
Spatial correspondence between effects of difficulty in semantic tasks and the top three dimensions of intrinsic connectivity. A, B and C – The first three connectivity dimensions identified through decomposition of the whole brain FC matrix. The first dimension corresponds to the principal gradient that separates sensory-motor regions (purple-blue) from transmodal areas (red-white). The second dimension separates auditory-motor cortex (purple-blue) from visual cortex (red-white). The third dimension separates FPCN regions (purple-blue) from DMN regions (red-white). D – The scree plot showing eigenvalue of each dimension. E and F – Unthresholded maps of the effects of difficulty in the semantic association and semantic feature matching tasks. G – Correlation between unthresholded effects of difficulty in each semantic task and the three connectivity dimensions. Effects of difficulty in the two semantic tasks dissociate within the brain space delineated by the dimensions of intrinsic connectivity, with effects of associative strength relating more to dimension 1, and effects of feature similarity relating more to dimension 3.

To elucidate the relationship between task difficulty effects of semantic tasks and the three connectivity dimensions, we calculated their spatial correlation across all brain parcels. All p-values were computed using spin permutation, which accounts for spatial autocorrelation, and were FDR corrected to control for multiple comparison. In the semantic association task, the difficulty effect positively correlated with the first dimension in the left hemisphere; control of the retrieval of global associations fell towards the heteromodal end of this component (LH: r = 0.32, p = 0.04; RH: r = 0.24, p = 0.09). There was no significant correlation with the second dimension, indicating a balanced recruitment of auditory-motor and visual processes during controlled retrieval of global associations (LH: r = 0.06, p = 0.39; RH: r = 0.02, p = 0.46). There was no significant correlation with the third dimension, suggesting an equal recruitment of control and DMN networks (LH: r = 0.04, p = 0.40; RH: r = 0.13, p = 0.16).

In contrast, the difficulty effect in the feature matching task negatively correlated with the first dimension in the right hemisphere, indicating difficulty modulated activation more in sensory-motor areas than heteromodal areas (LH: r = -0.24, p = 0.12; RH: r = - 0.36, p = 0.03). There was no correlation with the second dimension (LH: r = 0.36, p = 0.08; RH: r = 0.36, p = 0.08). However, a positive correlation was observed with the third dimension, showing stronger difficulty effects towards the control end than the DMN end (LH: r = 0.46, p = 0; RH: r = -0.33, p = 0.005).

Next, we compared the difficulty effects of the two semantic tasks within the brain state space. We calculated and transformed Pearson r correlations, which indicated the similarity between each connectivity dimension and the difficulty effect for each participant, to Fisher’s z values. The first dimension (heteromodal-unimodal) showed a stronger correlation with the effect of difficulty in semantic association task than feature matching task (t (27) = 3.921, p = 0.001; Fig 5G). This suggests that controlled retrieval in the association task more heavily involved heteromodal processes, whereas in the feature matching task, it was more modality-specific. The second dimension (visual-motor) had a stronger correlation with the effect of difficulty in feature matching than in semantic association (t (27) = -0.154, p = 0.019; Fig 5G), indicating that controlled responses in feature matching predominantly involved visual processing, while the association task employed a more balanced involvement of visual and motor information. Lastly, the third dimension (control-DMN) showed a greater correlation with the difficulty effect in feature matching than in association judgments (t (27) = -4.162, p = 0; Fig 5G). This indicates that feature matching relied more on the functional separation between domain-general executive processes and the long-term memory functions of the DMN, whereas the semantic association task engaged these networks in a more integrated manner (cf. Davey et al., 2016; Wang et al., 2020).

### 2.5. Comparison of the locations of difficulty effects in state space for semantic and non-semantic tasks

To compare the locations of difficulty effects in state space for semantic and non-semantic tasks, we first calculated correlations between non-semantic difficulty effects and the three dimensions. Fig. 6A and 6B show unthresholded difficulty effects for spatial working memory and math tasks, respectively. These spatial patterns correlated positively with the third dimension of intrinsic connectivity, which distinguishes control from DMN (spatial working memory - LH: r = 0.56, p = 0; RH: r = 0.60, p = 0; math tasks - LH: r = 0.61, p = 0; RH: r = 0.63, p = 0). There were no significant correlations with dimension 1 and 2 (uncorrected p > 0.05).

**Fig. 6.**
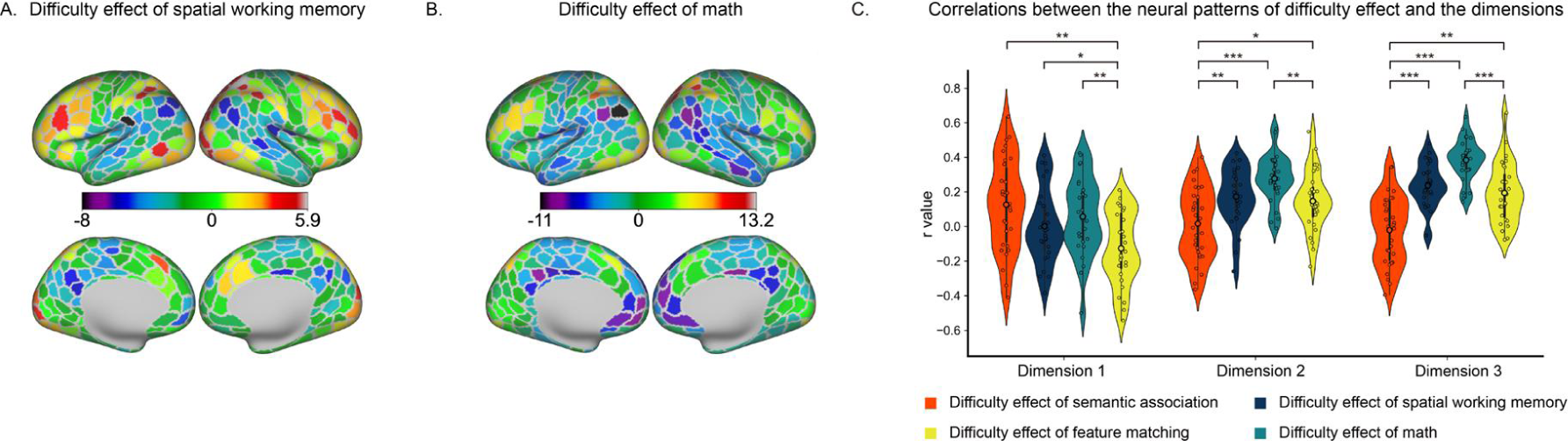
The spatial correspondence between effects of difficulty in non-semantic tasks and the dimensions of intrinsic connectivity. A and B – Unthresholded maps of the effects of difficulty in the spatial working memory and math tasks. C – The correlation between unthresholded effects of difficulty in each task and the three connectivity dimensions. Only the third dimension (control-DMN) correlated with the effects of difficulty in the two non-semantic tasks. The non-semantic tasks were also more similar to the feature matching than the association task on this dimension.

We then compared the correlations between connectivity dimensions and difficulty effects in the non-semantic tasks with the correlations between connectivity dimensions and difficulty effects in the semantic tasks. The first dimension of intrinsic connectivity was more associated with non-semantic difficulty than with task demands in the feature matching task (comparison for spatial working memory: t (26) = 2.26, p = 0.04; comparison for math: t (26) = 3.31, p = 0.006; Fig 6C). There were no differences between non-semantic difficulty and task demands in semantic association (spatial working memory: t (26) = -1.96, p = 0.07; math: t (26) = -1.12, p = 0.31; Fig 6C). These findings indicate that both semantic and non-semantic difficulty effects can fall towards the heteromodal end of the first dimension; in contrast, the feature matching task that involved the goal-driven retrieval of visual features for words was less heteromodal.

The second dimension of intrinsic connectivity, distinguishing visual from auditory-motor processes, showed greater correlation with non-semantic difficulty than task demands in the association matching task (spatial working memory versus association: t (26) = 3.09, p = 0.006; math versus association: t (26) = 5.467, p < 0.0001; Fig 6C). These results suggest that non-semantic tasks may involve more visual processing. Conversely, there was no significant difference between difficulty effects in spatial working memory and semantic feature matching (t (26) = 0.515, p = 0.609; Fig 6C); however, difficulty effects in the math task showed a stronger positive correlation than task demands in feature matching (t (26) = 2.963, p = 0.008; Fig 6C).

The third dimension of intrinsic connectivity, which separates control from DMN regions, correlated more strongly with difficulty effects in math compared with both semantic association (t (26) = 9.17, p < 0.0001; Fig 6C) and feature matching tasks (t (26) = 4.48, p < 0.0001; Fig 6C). Additionally, this dimension was more strongly correlated with spatial working memory than with task demands in semantic association (t (26) = 5.67, p = 0; Fig 5I), but no significant difference was found for feature matching (t (26) = 0.86, p = 0.39; Fig 5I). All the p-values were FDR corrected. These findings suggest that, on a dimension distinguishing control from DMN, difficulty effects in non-semantic tasks bear more similarity to those for feature matching than for global semantic associations.

## 3. Discussion

This study examines how cognitive control processes are organized on the cortical surface and within a brain state space defined by key dimensions of whole-brain intrinsic connectivity. We contrasted two semantic tasks — global association judgements and feature matching — and parametrically varied their difficulty by manipulating strength of association and feature similarity, to establish how brain networks are configured appropriately to control retrieval in these two contexts. We also compared controlled semantic cognition with the neural response to non-semantic control demands. We found that demanding semantic association trials elicited more activation in anterior portions of prefrontal and temporal cortex, while difficult semantic feature matching trials produced more posterior activation that overlapped to a greater extent with non-semantic multiple-demand regions. Differences were also found in whole-brain state space: the difficulty effects in global semantic associations were closer to the heteromodal end of a heteromodal-unimodal dimension than those in feature matching. Additionally, the association task demonstrated balanced recruitment between visual and auditory-motor representations on the second dimension and engaged both executive and DMN regions on the third dimension. In contrast, difficulty effects in semantic feature matching more closely resembled non-semantic task demands on the second and third dimensions, indicating greater visual and executive responses with less DMN involvement. These results collectively suggest there are at least two distinct large-scale brain states supporting controlled semantic cognition: one state is more heteromodal and involves more equal recruitment of control and DMN regions, while the other state is visually focused and engages control regions more selectively without concurrent DMN activation. Furthermore, these aspects control are underpinned by distinct dimensions of functional variation within whole-brain state space.

Semantic knowledge is multifaceted, drawing on support from diverse brain regions (Lambon Ralph et al., 2017). In our two semantic tasks, we utilized identical stimuli and presented them in the same format. Thus, the primary distinction between these tasks lies in the nature of the controlled retrieval process. The feature matching task predominantly relies on the controlled retrieval of visual features, while the semantic association task requires participants to draw upon heteromodal information since understanding the inherent relationships between word pairs involves integrating knowledge across various sensory experiences and modalities (Badre et al., 2005; Badre and Wagner, 2007; Gold et al., 2006). We show that the configuration of control processes that support cognition in a neural state space can reflect the type of information that participants are required to focus on, rather than simply the use of verbal materials, or the superficial characteristics of the task.

Recent research demonstrates that control regions modulate their activity and interaction patterns in a context-specific manner to support adaptable behavior across domains (Cole et al., 2013; Shine et al., 2019; Wang et al., 2023). These regions dynamically modify their baseline communication to integrate more specialized brain areas, facilitating task-specific computations (Finc et al., 2020; Khambhati et al., 2018; Koch et al., 2016). In neural state-space analysis, we found that this flexibility might relate to different network configurations underpinned by distinct dimensions of intrinsic connectivity. Specifically, control regions are proximal to DMN regions on the first dimension but are separated from DMN regions on the third dimension. This allows for whole-brain states in which heteromodal memory and control regions are either integrated (supporting task demands in association judgments) or segregated (supporting task demands in feature matching). These findings align with previous research suggesting that SCN and MDN are dissociable control networks: SCN appears to relate to the first neural dimension in which heteromodal memory and control networks are functionally coupled, while non-semantic controlled states linked to strong activation within MDN elicit anti-correlation between control and DMN regions, as captured by the third dimension (Jackson, 2021; Noonan et al., 2013; Wang et al., 2020, 2018). In line with this proposal, Zhang et al. (2021) found that regions of LIFG associated with maintaining and applying a semantic goal to constrain retrieval in a top-down fashion showed negative connectivity with DMN, while LIFG regions associated with the controlled retrieval of weak associations showed positive connectivity to some DMN regions. Neural state space analysis provides an account of both the commonalities and distinctions among various controlled states and explains why SCN and MDN are adjacent, yet topographically distinct.

Tasks involving global associations draw on diverse sensory-motor information, and therefore brain states that selectively focus on one modality are not conducive to the task. Here, control regions need to interact with heteromodal semantic knowledge to identify conceptual links between weakly related concepts and, consequently, heteromodal control and semantic memory networks are thought to be coupled in these circumstances (Davey et al., 2016). Consistent with this, control networks and DMN can show similar representational content (González-García et al., 2018; Wang et al., 2021) and both networks are modulated by prior knowledge (Gao et al., 2022; González-García et al., 2018). Conversely, tasks like visual feature matching demand a brain state in which visual (rather than auditory-motor) features dominate cognition. As decision-making hinges on one specific feature, control regions supporting goal maintenance and the prioritization of relevant knowledge need to be functionally separated from heteromodal conceptual knowledge and more tightly integrated with brain regions representing task-relevant information (Chiou and Lambon Ralph, 2016).

The concept of brain states offers a promising framework to understand neural flexibility and cognitive control, yet our study has limitations. Firstly, we focused on a neural state space defined by the top three dimensions of intrinsic connectivity, given these components explain the most variance and have clear interpretations in terms of functional relationships within and between heteromodal and unimodal cortex that are highly relevant to our task manipulations. However, cognitive control might be related to more than just these three dimensions. A more comprehensive understanding of the varieties of cognitive control will require exploring higher-dimensional state spaces. Secondly, although our tasks effectively demonstrate that distinct aspects of semantic control are related to different dimensions of brain state space, cognitive control can be modulated in numerous ways. Future research employing a broader array of tasks is essential to examine whether there are two primary dimensions of controlled behavior, one stabilized by heteromodal long-term memory and the other by control processes independent of memory. Despite these constraints, our study demonstrates that at least two neural dimensions are crucial to encompass the diverse range of controlled processes we employ to tailor cognition to the context.

## 4. Materials and Methods

### 4.1. Participants

All participants were right-handed, native English speakers, with normal or corrected-to-normal vision and no history of psychiatric or neurological illness. All participants provided informed consent. For the University of York datasets, the research was approved by the York Neuroimaging Centre and Department of Psychology ethics committees. For the HCP dataset, the study was approved by the Institutional Review Board of Washington University at St. Louis (Glasser et al., 2013).

31 healthy adults performed the semantic tasks (25 females; age: mean ± SD = 21.26 ± 2.93, range: 19 – 34 years). A functional run was excluded if (I) relative root mean square (RMS) framewise displacement was higher than 0.2 mm, (II) more than 15% of frames showed motion exceeding 0.25 mm, or (III) the accuracy of the behaviour task was low (3SD below the mean). If only one run of a task was left for a participant after exclusion, all their data for that task were removed. Using the exclusion criteria above for the feature matching task, there were 23 participants with 4 runs, 4 participants with 3 runs, and 1 participant with 2 runs. For the association task, there were 24 participants with 4 runs, 3 participants with 3 runs, and 3 participants with 2 runs. An additional 30 native English speakers, who did not take part in the main fMRI experiment, rated the color and shape similarity and semantic association strength for each word pair (21 females; age range: 18 – 24 years).

31 healthy adults (26 females; age: mean ± SD = 20.60 ± 1.68, range: 18 – 25 years) performed the spatial working memory and math tasks. One participant with incomplete data was removed. These exclusion criteria above resulted in a final sample of 27 participants for both the spatial working memory task and the math task.

The HCP sample involved data from 245 healthy volunteers (115 females), aged 23 – 35 years (mean = 28.21, SD = 3.67) (Glasser et al., 2013).

### 4.2. Task paradigms

#### 4.2.1. Semantic association task

Participants made yes/no decisions to pairs of words to indicate if they were semantically associated in general or not. Overall, there were roughly equal numbers of ‘related’ and ‘unrelated’ responses across participants. For example, DALMATIAN and COW are semantically related; COAL and TOOTH are not. Similarly, we parametrically manipulated the semantic association strength between the probe and target concepts, using semantic association strength ratings taken from a separate group of 30 participants on a 5-point Likert Scale. For example, in related trials, the association strength between PUMA and LION is very strong (i.e., 4.8) while for TIGER and WHALE is relatively weak (i.e., 4.0; although they are still both animals and are semantically related). In non-related trials, the association strength between KINGFISHER and SCORPION is relatively high (i.e., 2.1) while BANANA and BRICK is very low (i.e., 1.0) although participants thought neither were related. For the related trials, stronger associations would facilitate decision making, while for unrelated trials, stronger associations interfere with the decision making. This parametric design allowed us to model the effect of decision difficulty and test whether how this is related to dimensions of brain organization.

This task included four runs, presented in a rapid event-related design. Each run consisted of 80 trials, with about half being related and half being unrelated trials. The procedure was the same as the feature matching task except only two words were presented on the screen. The feature and association tasks were separated by one week.

#### 4.2.2. Semantic feature matching task

Participants made yes/no decisions about whether probe and target concepts (presented as words) were matched in terms of a particular semantic feature (colour or shape), specified at the top of the screen during each trial. The feature prompt, probe word, and target words were presented simultaneously. Half of the trials were matching trials in which participants were expected to identify shared features; while half of the trials were non-matching trials in which participants would not be expected to identify shared features. For example, in a colour matching trial, participants would answer ‘yes’ to the word-pair DALMATIAN – COW, due to their colour similarity, whereas they would answer ‘no’ to COAL – TOOTH as they do not share a similar colour. The same stimuli were used in the semantic feature matching task and semantic association task.

We parametrically manipulated the degree of feature similarity between the probe and target concepts, using semantic feature similarity ratings taken from a separate group of 30 participants on a 5-point Likert Scale. For instance, in colour-matching trials, the degree of colour similarity between DALMATIAN and COW was found to be very high (i.e., 4.8), while that between PUMA and LION was relatively low (i.e., 4.0), despite that participants believe that the two trials had similar colour. Conversely, in colour non-matching trials, the degree of colour similarity between CROW and HUMMINGBIRD was relatively high (i.e., 2.5), whereas that between COAL and TOOTH was very low (i.e., 1.0), even though the participants perceived no similarity in colour. Greater feature similarity facilitates the decision-making process for the matching trials but makes the decision more difficult for the non-matching trials. This parametric design allowed us to model the effect of the decision difficulty during the controlled retrieval of visual features in the neural data, and test how it is related to dimensions of brain organization.

This task included four runs and two conditions (two features: colour and shape), presented in a mixed design. Each run consisted of four experimental blocks (two 2 min 30 s blocks per feature), resulting in a total time of 10 min 12 s. In each block, 20 trials were presented in a rapid event-related design. To maximize the statistical power of the rapid event-related fMRI data analysis, the stimuli were presented with a temporal jitter randomized from trial to trial (Dale, 1999). The inter-trial interval varied from 3 to 5 s. Each trial started with a fixation, followed by the feature, probe word, and target word presented centrally on the screen, triggering the onset of the decision-making period. The feature, probe word, and target word remained visible until the participant responded, or for a maximum of 3 s. The condition order was counterbalanced across runs and run order was counterbalanced across participants. Half of the participants pressed a button with their right index finger to indicate a matching trial and responded with their right middle finger to indicate a non-matching trial. Half of the participants pressed the opposite buttons.

#### 4.2.3. Spatial working memory task

Participants were required to maintain four or eight sequentially presented locations in a 3×4 grid (Fedorenko et al., 2011), giving rise to easy and hard spatial working memory conditions. Stimuli were presented at the center of the screen across four steps. Each of these steps lasted for 1s and highlighted one location on the grid in the easy condition, and two locations in the hard condition. This was followed by a decision phase, which showed two grids side by side (i.e., two-alternative forced choice (2AFC) paradigm). One grid contained the locations shown on the previous four steps, while the other contained one or two locations in the wrong place. Participants indicated their response via a button press and feedback was immediately provided within in 2.5s.

Each run consisted of 12 experimental blocks (6 blocks per condition and 4 trials in a 32 s block) and 4 fixation blocks (each 16 s long), resulting in a total time of 448 s.

#### 4.2.4. Math task

Participants were presented with an addition expression on the screen for 1.45s and, subsequently made a 2AFC decision indicating their solution within 1s. The easy condition used single-digit numbers while the hard condition used two-digit numbers. Each trial ended with a blank screen lasting for 0.1s. Each run consisted of 12 experimental blocks (with 4 trials per block) and 4 fixation blocks, resulting in a total time of 316s.

### 4.3. Image acquisition

#### 4.3.1. Image acquisition of York Semantic dataset

Whole brain structural and functional MRI data were acquired using a 3T Siemens MRI scanner utilising a 64-channel head coil, tuned to 123 MHz at York Neuroimaging Centre, University of York. The functional runs were acquired using a multi-band multi-echo (MBME) EPI sequence, each 11.45 minutes long (TR=1.5 s; TE = 12, 24.83, 37.66 ms; 48 interleaved slices per volume with slice thickness of 3 mm (no slice gap); FoV = 24 cm (resolution matrix = 3x3x3; 80x80); 75° flip angle; 455 volumes per run; 7/8 partial Fourier encoding and GRAPPA (acceleration factor = 3, 36 ref. lines); multi-band acceleration factor = 2). Structural T1-weighted images were acquired using an MPRAGE sequence (TR = 2.3 s, TE = 2.3 s; voxel size = 1x1x1 isotropic; 176 slices; flip angle = 8°; FoV= 256 mm; interleaved slice ordering). We also collected a high-resolution T2- weighted (T2w) scan using an echo-planar imaging sequence (TR = 3.2 s, TE = 56 ms, flip angle = 120°; 176 slices, voxel size = 1x1x1 isotropic; Fov = 256 mm).

#### 4.3.2. Image acquisition of York Non-semantic dataset

MRI acquisition protocols have been described previously (Wang et al., 2021; Wang et al., 2020). Structural and functional data were collected on a Siemens Prisma 3T MRI scanner at the York Neuroimaging Centre. The scanning protocols included a T1- weighted MPRAGE sequence with whole-brain coverage. The structural scan used: acquisition matrix of 176 × 256 × 256 and voxel size 1 × 1 × 1 mm^3^, repetition time (TR) = 2300 ms, and echo time (TE) = 2.26 ms. Functional data were acquired using an EPI sequence with an 800 flip angle and using GRAPPA with an acceleration factor of 2 in 3 x 3 x 4 mm voxels in 64-axial slices. The functional scan used: 55 3-mm-thick slices acquired in an interleaved order (with 33% distance factor), TR = 3000 ms, TE = 15 ms, FoV = 192 mm.

#### 4.3.3. Image acquisition of HCP dataset

MRI acquisition protocols of the HCP dataset have been previously described (Barch et al., 2013; Glasser et al., 2013). Images were acquired using a customized 3T Siemens Connectome scanner having a 100 mT/m SC72 gradient set and using a standard Siemens 32-channel radiofrequency receive head coil. Participants underwent the following scans: structural (at least one T1-weighted (T1w) MPRAGE and one 3D T2- weighted (T2w) SPACE scan at 0.7-mm isotropic resolution), rsfMRI (4 runs ×14 min and 33 s), and task fMRI (7 tasks, 46.6 min in total). Since not all participants completed all scans, we only included 339 unrelated participants from the S900 release. Whole-brain rsfMRI and task fMRI data were acquired using identical multi-band echo planar imaging (EPI) sequence parameters of 2-mm isotropic resolution with a TR = 720 ms. Spin echo phase reversed images were acquired during the fMRI scanning sessions to enable accurate cross-modal registrations of the T2w and fMRI images to the T1w image in each subject and standard dual gradient echo field maps were acquired to correct T1w and T2w images for readout distortion. Additionally, the spin echo field maps acquired during the fMRI session (with matched geometry and echo spacing to the gradient echo fMRI data) were used to compute a more accurate fMRI bias field correction and to segment regions of gradient echo signal loss.

Subjects were considered for data exclusion based on the mean and mean absolute deviation of the relative root-mean-square motion across four rsfMRI scans, resulting in four summary motion measures. If a subject exceeded 1.5 times the interquartile range (in the adverse direction) of the measurement distribution in two or more of these measures, the subject was excluded. In addition, functional runs were flagged for exclusion if more than 25% of frames exceeded 0.2 mm frame-wise displacement (FD_power). These above exclusion criteria were established before performing the analysis (Faskowitz et al., 2020; Sporns et al., 2021). The data of 91 participants was excluded because of excessive head motion and the data of another 3 participants was excluded because their resting data did not have all the time points. In total, the data of 245 participants was analysed after exclusions.

### 4.4. Image pre-processing

#### 4.4.1. Image pre-processing of York Semantic and Non-semantic dataset

The York datasets were preprocessed using fMRIPrep 20.2.1 [(Esteban et al., 2018), RRID:SCR_016216], which is based on Nipype 1.5.1 [(Gorgolewski et al., 2011), RRID:SCR_002502].

##### 4.4.1.1. Anatomical data preprocessing

The T1w image was corrected for intensity non-uniformity (INU) with N4BiasFieldCorrection (Tustison et al., 2010), distributed with ANTs 2.3.3 [(Avants et al., 2008), RRID:SCR_004757], and used as T1w-reference throughout the workflow. The T1w-reference was then skull-stripped with a Nipype implementation of the antsBrainExtraction.sh workflow (from ANTs), using OASIS30ANTs as target template. Brain tissue segmentation of cerebrospinal fluid (CSF), white-matter (WM) and gray-matter (GM) was performed on the brain-extracted T1w using fast FSL 5.0.9 [(Zhang et al., 2001), RRID:SCR_002823]. Brain surfaces were reconstructed using recon-all from FreeSurfer 6.0.1 [(Dale et al., 1999a), RRID:SCR_001847], and the brain mask estimated previously was refined with a custom variation of the method to reconcile ANTs-derived and FreeSurfer-derived segmentations of the cortical gray-matter of Mindboggle [(Klein et al., 2017), RRID:SCR_002438]. Volume-based spatial normalization to two standard spaces (MNI152NLin2009cAsym, MNI152NLin6Asym) was performed through nonlinear registration with antsRegistration (ANTs 2.3.3), using brain-extracted versions of both T1w reference and the T1w template. The following templates were selected for spatial normalization: ICBM 152 Nonlinear Asymmetrical template version 2009c [(Fonov et al., 2009), RRID:SCR_008796; TemplateFlow ID: MNI152NLin2009cAsym], FSL’s MNI ICBM 152 non-linear 6th Generation Asymmetric Average Brain Stereotaxic Registration Model [(Evans et al., 2012), RRID:SCR_002823; TemplateFlow ID: MNI152NLin6Asym].

##### 4.4.1.2. Functional data preprocessing

For each of the BOLD runs per subject, the following preprocessing was performed. First, a reference volume and its skull-stripped version were generated using a custom methodology of fMRIPrep. A B0-nonuniformity map (or fieldmap) was estimated based on a phase-difference map calculated with a dual-echo GRE (gradient-recall echo) sequence, processed with a custom workflow of SDCFlows inspired by the epidewarp.fsl script and further improvements in HCP Pipelines (Glasser et al., 2013). The fieldmap was then co-registered to the target EPI reference run and converted to a displacements field map (amenable to registration tools such as ANTs) with FSL’s fugue and other SDCflows tools. Based on the estimated susceptibility distortion, a corrected EPI reference was calculated for a more accurate co-registration with the anatomical reference. The BOLD reference was then co-registered to the T1w reference using bbregister (FreeSurfer) which implements boundary-based registration (Greve and Fischl, 2009). Co-registration was configured with six degrees of freedom. Head-motion parameters with respect to the BOLD reference (transformation matrices, and six corresponding rotation and translation parameters) were estimated before any spatiotemporal filtering using mcflirt (FSL 5.0.9, (Jenkinson et al., 2002)). BOLD runs were slice-time corrected using 3dTshift from AFNI 20160207 [(27), RRID:SCR_005927]. The BOLD time-series were resampled onto the following surfaces (FreeSurfer reconstruction nomenclature): fsaverage. Grayordinates files (Glasser et al., 2013) containing 91k samples were also generated using the highest-resolution fsaverage as intermediate standardized surface space. Several confounding time-series were calculated based on the preprocessed BOLD: framewise displacement (FD), DVARS (D refers to a derivative of fMRI time course, VARS refers to RMS variance) and three region-wise global signals. FD was computed using two formulations following previous work (absolute sum of relative motion; (Power et al., 2014), relative root mean square displacement between affines; (Jenkinson et al., 2002). FD and DVARS were calculated for each functional run, both using their implementations in Nipype (Power et al., 2014). Three global signals were extracted within the CSF, the WM, and the whole- brain masks. Additionally, a set of physiological regressors were extracted to allow for component-based noise correction (CompCor) (Behzadi et al., 2007) principal components were estimated after high-pass filtering the preprocessed BOLD time-series (using a discrete cosine filter with 128s cut-off) for two CompCor variants: temporal (tCompCor) and anatomical (aCompCor). tCompCor components were then calculated from the top 2% variable voxels within the brain mask. For aCompCor, three probabilistic masks (CSF, WM and combined CSF+WM) were generated in anatomical space. The implementation differs from that of Behzadi et al. (Behzadi et al., 2007) in that instead of eroding the masks by 2 pixels in BOLD space, the aCompCor masks are subtracted from a mask of pixels that likely contain a volume fraction of GM. This mask is obtained by dilating a GM mask extracted from the FreeSurfer’s aseg segmentation, and it ensures components are not extracted from voxels containing a minimal fraction of GM. Finally, these masks are resampled into BOLD space and binarized by thresholding at 0.99 (as in the original implementation). Components were also calculated separately within the WM and CSF masks. For each CompCor decomposition, the k components with the largest singular values were retained, such that the retained components’ time series were sufficient to explain 50 percent of variance across the nuisance mask (CSF, WM, combined, or temporal). The remaining components were dropped from consideration. The head-motion estimates calculated in the correction step were also placed within the corresponding confounds file. The confound time series derived from head motion estimates and global signals were expanded with the inclusion of temporal derivatives and quadratic terms for each (Satterthwaite et al., 2013). Frames that exceeded a threshold of 0.5 mm FD or 1.5 standardized DVARS were annotated as motion outliers. All resamplings were performed with a single interpolation step by composing all the pertinent transformations (i.e., head-motion transform matrices, susceptibility distortion correction when available, and co-registrations to anatomical and output spaces). Gridded (volumetric) resamplings were performed using antsApplyTransforms (ANTs), configured with Lanczos interpolation to minimize the smoothing effects of other kernels (Lanczos, 1964). Non-gridded (surface) resamplings were performed using mri_vol2surf (FreeSurfer). fMRIPrep used Nilearn 0.6.2 [(Abraham et al., 2014) RRID:SCR_001362], mostly within the functional processing workflow. The resulting data were in CIFTI 64k- vertex grayordinate space. The left hemisphere had 29696 vertices and right hemisphere had 29716 vertices in total after removing the medial wall.

Post-processing of the outputs of fMRIPrep version 20.2.1 (Esteban et al., 2018) was performed using the eXtensible Connectivity Pipeline (XCP) (Satterthwaite et al., 2013; Ciric et al., 2018). For each CIFTI run per subject, the following post-processing was performed: before nuisance regression and filtering any volumes with framewise-displacement greater than 0.3 mm (Satterthwaite et al., 2013; Power et al., 2014) were flagged as outliers and excluded from nuisance regression. In total, 36 nuisance regressors were selected from the nuisance confound matrices of fMRIPrep output. These nuisance regressors included six motion parameters, global signal, mean white matter, and mean CSF signal with their temporal derivatives, and the quadratic expansion of six motion parameters, tissue signals and their temporal derivatives (Satterthwaite et al., 2013; Ciric et al., 2017, 2018). These nuisance variables were accounted for in the BOLD data using linear regression - as implemented in Scikit-Learn 0.24.2 (Pedregosa et al., 2011). Residual timeseries from this regression were then band-pass filtered to retain signals within the 0.01-0.08 Hz frequency band. The processed BOLD was smoothed using Connectome Workbench with a gaussian kernel size of 6.0 mm (FWHM). Processed functional timeseries were extracted from residual BOLD using Connectome Workbench (Glasser et al., 2013) for the Glasser atlas (Glasser et al., 2016). Many internal operations of XCP use Nibabel (Abraham et al., 2014), numpy (Harris et al., 2020), and scipy (Harris et al., 2020).

#### 4.4.2. Image pre-processing of HCP dataset

We used HCP’s minimal pre-processing pipelines (Glasser et al., 2013). Briefly, for each subject, structural images (T1w and T2w) were corrected for spatial distortions. FreeSurfer v5.3 was used for accurate extraction of cortical surfaces and segmentation of subcortical structures (Dale et al., 1999b; Fischl et al., 1999). To align subcortical structures across subjects, structural images were registered using non-linear volume registration to the Montreal Neurological Institute (MNI152) space. Functional images (rest and task) were corrected for spatial distortions, head motion, and mapped from volume to surface space using ribbon-constrained volume to surface mapping.

Subcortical data were also projected to the set of extracted subcortical structure voxels and combined with the surface data to form the standard CIFTI grayordinate space. Data were smoothed by a 2-mm FWHM kernel in the grayordinates space that avoids mixing data across gyral banks for surface data and avoids mixing areal borders for subcortical data. Rest and task fMRI data were additionally identically cleaned for spatially specific noise using spatial ICA+FIX (Salimi-Khorshidi et al., 2014) and global structured noise using temporal ICA (Glasser et al., 2018). For accurate cross-subject registration of cortical surfaces, a multimodal surface matching (MSM) algorithm (Robinson et al., 2014) was used to optimize the alignment of cortical areas based on features from different modalities. MSMSulc (“sulc”: cortical folds average convexity) was used to initialize MSMAll, which then utilized myelin, resting-state network, and rfMRI visuotopic maps.

### 4.5. Task fMRI analysis

#### 4.5.1. Individual-specific parcellation

Considering the anatomical and functional variability across individuals (Braga and Buckner, 2017; Gordon et al., 2017; Laumann et al., 2015; Mueller et al., 2013), we estimated individual-specific areal-level parcellation using a multi-session hierarchical Bayesian model (MS-HBM) (Kong et al., 2021, 2019). To estimate individual-specific parcellation, we acquired ‘‘pseudo-resting state’’ timeseries in which the task activation model was regressed from feature matching and semantic association fMRI data (Fair et al., 2007) using xcp_d (https://github.com/PennLINC/xcp_d). The task activation model and nuisance matrix were regressed out using AFNI’s3dTproject (for similar implementation, see Cui et al. (2020).

Using a group atlas, this method calculates inter-subject resting-state functional connectivity variability, intra-subject resting-state functional connectivity variability, and finally parcellates for each single subject based on this prior information. As in Kong et al. (Kong et al., 2021, 2019), we used MS-HBM to define 400 individualized parcels belonging to 17 discrete individualized networks for each participant. Specifically, we calculated all participants’ connectivity profiles, created the group parcellation using the average connectivity profile of all participants, estimated the inter-subject and intra-subject connectivity variability, and finally calculated each participant’s individualized parcellation. This parcellation imposed the Markov random filed (MRF) spatial prior. We used a well-known areal-level parcellation approach, i.e., the local gradient approach (gMS-HBM), which detects local abrupt changes (i.e., gradients) in resting-state functional connectivity across the cortex (Cohen et al., 2008). A previous study (Schaefer et al., 2018) has suggested combining local gradient (Cohen et al., 2008; Gordon et al., 2016) and global clustering (Yeo et al., 2011) approaches for estimating areal-level parcellations. Therefore, we complemented the spatial contiguity prior in contiguous MS-HBM (cMS-HBM) with a prior based on local gradients in resting-state functional connectivity, which encouraged adjacent brain locations with gentle changes in functional connectivity to be grouped into the same parcel. We used the pair of parameters (i.e., beta value = 50, w = 30 and c = 30), which was optimized using our own dataset. The same parameters were also used in Kong et al. (Kong et al., 2021). Vertices were parcellated into 400 cortical regions (200 per hemisphere). To parcellate each of these parcels, we calculated the average time series of enclosed vertices to get better signal noise ratio (SNR) using Connectome Workbench software. This parcel-based time series was used for all the following analyses. The same method and parameters were used to generate the individual-specific parcellation for the participants in the HCP dataset using the resting-state time series except that the task regression was not performed.

##### 4.5.1.1 Homogeneity of parcels

To evaluate whether a functional parcellation is successful, parcel homogeneity is commonly used (Gordon et al., 2016; Kong et al., 2019, 2021). Parcel homogeneity was calculated as the average Pearson’s correlations between fMRI time courses of all pairs of vertices within each parcel, adjusted for parcel size and summed across parcels (Schaefer et al., 2018; Kong et al., 2019, 2021). Higher homogeneity means that vertices within the same parcel share more similar time courses and indicates better parcellation quality. To summarize the parcel homogeneity, we averaged the homogeneity value across parcels. We calculated the parcel homogeneity for each run of each participant for each task using the individual-specific parcellation and then averaged them across runs for each participant for each task. We also calculated the parcel homogeneity using canonical Yeo 17-network group atlas. Using the resting state data of the HCP dataset, Kong et al. (2021) demonstrated that homogeneity within MS-HBM-based individualized parcels was greater than that in the canonical Yeo 17-network group atlas that does not consider variation in functional neuroanatomy. A similar pattern was observed using the York Semantic datasets (Wang et al., 2023).

#### 4.5.2. Task fMRI univariate analysis

To reveal how the neural data were modulated by the difficulty of making decisions about global semantic associations and visual features, respectively, we conducted univariate analysis for the association task and feature matching task, respectively and then compared them. To examine parametric effects of task difficulty, we modelled the parametric effect of associative strength or feature similarity, including a parametric regressor for correct trials in the general linear model (GLM). Additionally, we included one task mean regressor to reveal the main effect of task, which is analogous to the inclusion of an intercept term in a linear regression model along with the slope term. The task mean effect was used to reveal the regions that showed greater or less activation during the tasks relative to the rest by extracting the beta value of each parcel in these task conditions and testing whether they were significantly activated (i.e., above zero) or deactivated (i.e., below zero) relative to implicit baseline (i.e., fixation period). For all the tasks, we also modelled incorrect trials as regressors of no interest. Demeaned semantic ratings and the main effect of task were modelled as epochs lasting from the trial onset to response, thus controlling for lengthened BOLD responses on trials with longer response times. Fixed-effects analyses were conducted using nilearn (Abraham et al., 2014) to estimate the average effects across runs within each subject for each parcel. Then we conducted one-sample t-tests to assess whether the estimated effect-size (i.e., contrast) was significantly different from zero across all subjects. We conducted FDR correction at p = 0.05 to control for multiple comparisons. Finally, we identified the network that each parcel belonged to (Kong et al., 2021).

Then, we examined the difficulty effect for each task. We pinpointed brain regions that exhibited a stronger response to more difficult trials in the two semantic tasks. This increase in activation occurred when (i) association strength was lower for related ’Yes’ trials or higher for unrelated ’No’ trials in the semantic association task, and (ii) feature similarity was lower for matching ’Yes’ trials or higher for non-matching ’No’ trials in the feature matching task.

In the semantic association task, we modeled the parametric effect of difficulty using demeaned semantic association strength ratings. Our analysis focused on how neural responses varied with association strength: they were negatively modulated by association strength in related trials and positively modulated in non-related trials. Additionally, we identified brain regions that exhibited increased activation during easier trials, characterized by comparatively weak associative strength in associated trials and strong associative strength in non-associated trials.

Similarly, we examined the difficulty effect for the semantic feature matching task. We modeled the difficulty effect using demeaned feature similarity ratings. We examined how neural responses were modulated by these ratings: they were negatively modulated by feature similarity in matching trials and positively in non-matching trials. To identify specific brain regions involved, we extracted the beta values for each parcel. This helped reveal regions that demonstrated greater activation when feature similarity was lower in matching trials and higher in non-matching trials. Additionally, we identified regions that showed the opposite pattern, exhibiting greater deactivation in easier trials (i.e., when feature similarity was lower in matching trials and higher in non-matching trials). To directly compare differences in the activation patterns for the association judgment and feature matching tasks, we extracted the beta values relating to semantic difficulty for each parcel and each participant in each task and conducted paired t-tests.

We also examined regions where the neural responses were modulated by task difficulty in spatial working memory and math tasks. We included two regressors – hard and easy conditions to reveal regions showing greater activation in the hard than easy conditions. These parcels were thought to support domain-general executive control. We also modelled incorrect trials as regressors of no interest.

#### 4.5.3. Comparison of semantic and non-semantic task demands

After determining the difficulty effects of both semantic and non-semantic tasks, we analyzed the extent of overlap between these effects in semantic tasks and brain regions responsive to non-semantic task demands through three complementary analyses. Firstly, we quantified the overlap in regions showing greater activation in semantic association task with those in either spatial working memory or math tasks. We also quantified such overlap for the semantic feature matching task. Secondly, we identified MDN regions by locating areas with difficulty effects in both spatial working memory and math tasks. We then compared the activation strength linked to task difficulty in these MDN regions for both semantic association and feature matching tasks. Lastly, we calculated and compared spatial correlations between the unthresholded maps of difficulty effects in non-semantic and semantic tasks. These analyses enabled us to investigate if the difficulty effect in the feature matching task showed a greater overlap with non-semantic control areas compared to the semantic association task. All p-values were FDR-corrected following spin permutation.

Given the spatial autocorrelation present in the task difficulty maps, we created a null distribution using spin permutation implemented in BrainSMASH (Burt et al., 2020). This approach simulates brain maps, constrained by empirical data, that preserve the spatial autocorrelation of cortical parcellated brain maps. We subsequently compared the observed correlation values with the null distribution to determine whether the real correlations were significantly greater than that expected by spatial autocorrelation alone. This analysis was performed for the two hemispheres separately because the geodesic distance between parcels was used to generate the spatial-autocorrelation-preserving surrogate maps when creating the null distribution, and we could only measure geodesic distance between parcels within a hemisphere, because the left and right hemisphere surface maps were not on the same mesh.

#### 4.5.4. The dimensions of intrinsic connectivity

We identified key dimensions of FC by performing dimension reduction analysis on resting state FC from the HCP dataset. First, we calculated the resting-state functional connectivity for each run of each participant by demeaning the residual time series for each parcel and then calculating the Pearson correlations for each parcel pair. We then averaged these individual connectivity matrices to generate a group-averaged connectivity matrix. We used the Brainspace Toolbox (Vos de Wael et al., 2020) to extract ten group-level gradients from the group-averaged connectivity matrix (dimension reduction technique = diffusion embedding, kernel = None, sparsity = 0.9), following the methodology of previous studies (Mckeown et al., 2020; Wang et al., 2020). This analysis resulted in ten group-level gradients explaining maximal whole-brain connectivity variance in descending order. We retained the first few components explaining the most variance by looking at the eigenvalues of each component in the scree plots shown in Fig 5D. The first three components, which explained 28.02% variance, had the largest eigenvalues, indicating their greater importance (see Fig. 5D for scree plot)

#### 4.5.5. Correlation between parametric difficulty effects and connectivity components

We investigated whether the primary dimensions of brain organization, as captured by connectivity components, correspond to the topographical organization of the parametric effects of task difficulty. The semantic association task may rely more on the separation between sensory-motor and transmodal regions, essential for the controlled retrieval of long-term memory. Conversely, the feature matching task may rely more on the separation between domain-general control network and DMN, due to its goal maintenance demands that typically engage control networks that are anti-correlated with DMN. We examined the relationship between task difficulty effects, indicated by parametric regressors, and functional organization dimensions, revealed through intrinsic connectivity components. This involved computing Pearson r correlations between the first three connectivity dimensions and difficulty effects of semantic and non-semantic tasks at the group level. Given the spatial autocorrelation present in both the principal connectivity gradient and task difficulty maps, we created a null distribution using spin permutation implemented in BrainSMASH (Burt et al., 2020).

To compare the locations of difficulty effects in state space for semantic and non-semantic tasks, we also calculated the Pearson r correlation between the first three connectivity components and the difficulty effect for each task for each participant and then converted the Pearson r values to Fisher z values. Finally, we compared the correlations for each task pair by conducting paired-t test. We conducted FDR correction at p = 0.05 to control for multiple comparisons.

### 4.6. Structural MRI analysis

#### 4.6.1. Cortical geometry - global minimum distance to primary sensory-motor landmarks

We investigated whether the demanding semantic association task elicited more anterior brain responses, located further from the sensory-motor cortex, compared to the semantic feature matching task. To do this, we analyzed how closely these responses were located to the sensory-motor cortex. Specifically, we classified brain parcels into four groups according to their response to task difficulty: (i) parcels responding only during the semantic association task, (ii) parcels showing responses in both semantic association and non-semantic tasks, (iii) parcels affected in both feature matching and non-semantic tasks, and (iv) parcels responsive exclusively during the feature matching task. We then calculated the shortest distance (global minimum distance) from each parcel to the nearest sensory-motor landmarks for each participant.

We calculated the geodesic distance between each parcel and key landmarks associated with primary visual, auditory and somatomotor cortex. These values were used to identify the minimum geodesic distance to primary sensory-motor regions for each parcel. Three topographical landmarks were used: the central sulcus corresponding to the primary somatosensory/motor cortex; temporal transverse sulcus indicating primary auditory cortex; and calcarine sulcus, marking the location of primary visual cortex. Since the cortical folding patterns vary across participants, and the individual variability in cortical folding increases with cortical surface area (Van Essen et al., 2019), both the shapes of these landmarks and the number of vertices within each landmark might show individual differences. We used participant-specific landmark label files to locate the participant-specific vertices belonging to each landmark and participant-specific parcellation to locate the vertices within each parcel.

Geodesic distance along the ‘midthickness’ of the cortical surface (halfway between the pial and white matter) was calculated using the Connectome Workbench software with an algorithm that measures the shortest path between two vertices on a triangular surface mesh (Mitchell et al., 1987; O’Rourke, 1999). This method returns distance values independent of mesh density. Geodesic distance was extracted from surface geometry (GIFTI) files, following surface-based registration (Robinson et al., 2014). To ensure that the shortest paths would only pass through the cortex, vertices representing the medial wall were removed from the triangular mesh for this analysis.

We calculated the minimum geodesic distance between each vertex and each landmark. Specifically, for the central sulcus, we calculated the geodesic distance between vertex i outside the central sulcus and each vertex within it (defined for each individual). We then identified vertex j within the central sulcus closest to vertex i, and extracted this value as the minimum geodesic distance for vertex i to this landmark. To compute the minimum geodesic distance for parcel k to the central sulcus, we averaged the minimum distance across all the grayordinate vertices in parcel k to the vertices within the central sulcus. The same procedure was applied to calculate minimum geodesic distance between each parcel and all three sensory-motor landmarks (central sulcus, temporal transverse sulci, and calcarine sulcus). From these three minimum geodesic distances, we selected the lowest distance value (i.e., the closest landmark to parcel k) as the global minimum distance to sensory-motor regions for parcel k. Then we averaged the mean minimum distance of all the parcels within each type of parcels for each participant. Finally, we examined whether mean minimum distance of each type of parcels were different by performing a paired t-test. All p-values are FDR-corrected.

### 4.7. Data and Code availability

The HCP data is publicly available here https://www.humanconnectome.org/. The York data is not available due to insufficient consent. Researchers wishing to access the data should contact Elizabeth Jefferies or the Chair of the Research Ethics Committee of the York Neuroimaging Centre. Data will be released when this is possible under the terms of the UK GDPR. Analysis code for this study is available at https://github.com/Xiuyi-Wang/Project_Semantic_Gradient.

## Supporting information

Macroscale brain states support the control of semantic cognition

## Author contributions

X.W., E.J. designed research; K.K.R., and X.W. collected the data, X.W. analyzed data; X.W., Y.D., and E.J. wrote the original manuscript. All authors edited the manuscript.

## Acknowledgements

We are grateful to Pradeepa Ruwan and Antonia De Freitas for piloting the experiment. X. W. discloses support for the research of this work from Scientific Foundation of Institute of Psychology, Chinese Academy of Sciences (Grant No. E1CX4725CX) and the National Natural Science Foundation of China (Grant No. 32300881). Y.D. discloses support for the publication of this work from the STI 2030—Major Projects (Grant Number. 2021ZD0201500), the National Natural Science Foundation of China (Grant No. 31822024), and Scientific Foundation of Institute of Psychology, Chinese Academy of Sciences (Grant Number. E2CX3625CX). The research was supported by a European Research Council Consolidator grant (Project ID: 771863 - FLEXSEM) to E.J.

