## Supplementary material for "Macroscale brain states support the control of semantic cognition": Macroscale brain states support the control of semantic cognition

### Author contributions

X.W., E.J. designed research; K.K.R., and X.W. collected the data, X.W. analyzed data; X.W., Y.D., and E.J. wrote the original manuscript. All authors edited the manuscript.

### Acknowledgements

We are grateful to Pradeepa Ruwan and Antonia De Freitas for piloting the experiment. X. W. discloses support for the research of this work from Scientific Foundation of Institute of Psychology, Chinese Academy of Sciences (Grant No. E1CX4725CX) and the National Natural Science Foundation of China (Grant No. 32300881). Y.D. discloses support for the publication of this work from the STI 2030—Major Projects (Grant Number. 2021ZD0201500), the National Natural Science Foundation of China (Grant No. 31822024), and Scientific Foundation of Institute of Psychology, Chinese Academy of Sciences (Grant Number. E2CX3625CX). The research was supported by a European Research Council Consolidator grant (Project ID: 771863 - FLEXSEM) to E.J.

### Declaration of interests

The authors declare no competing interests.

### Supplementary Materials

##### 1.1. Semantic association and feature matching judgements showed similar task activation and deactivation pattern

We established that the semantic association and semantic feature matching tasks evoked similar patterns of activation and deactivation. We identified these regions by constructing a general linear model (GLM) and comparing tasks to rest, including both Yes and No trials. As expected, multiple regions in the DAN and FPCN, including inferior parietal lobe, inferior parietal sulcus and superior frontal gyrus etc., exhibited activation, while regions in the DMN, including angular gyrus, anterior temporal lobe, posterior cingulate cortex etc., showed deactivation during both semantic association (Fig. S1A) and feature matching (Fig. S1B). 62% of regions showing BOLD increases in either task were activated by both tasks, and 72% of regions deactivating in either task showed common deactivation across two tasks (Fig. S1C). A few regions in the visual network showed stronger activation during semantic association than semantic feature matching task, while a few regions in motor, DAN, FPCN-A networks demonstrated the reverse pattern (Fig. S1D).

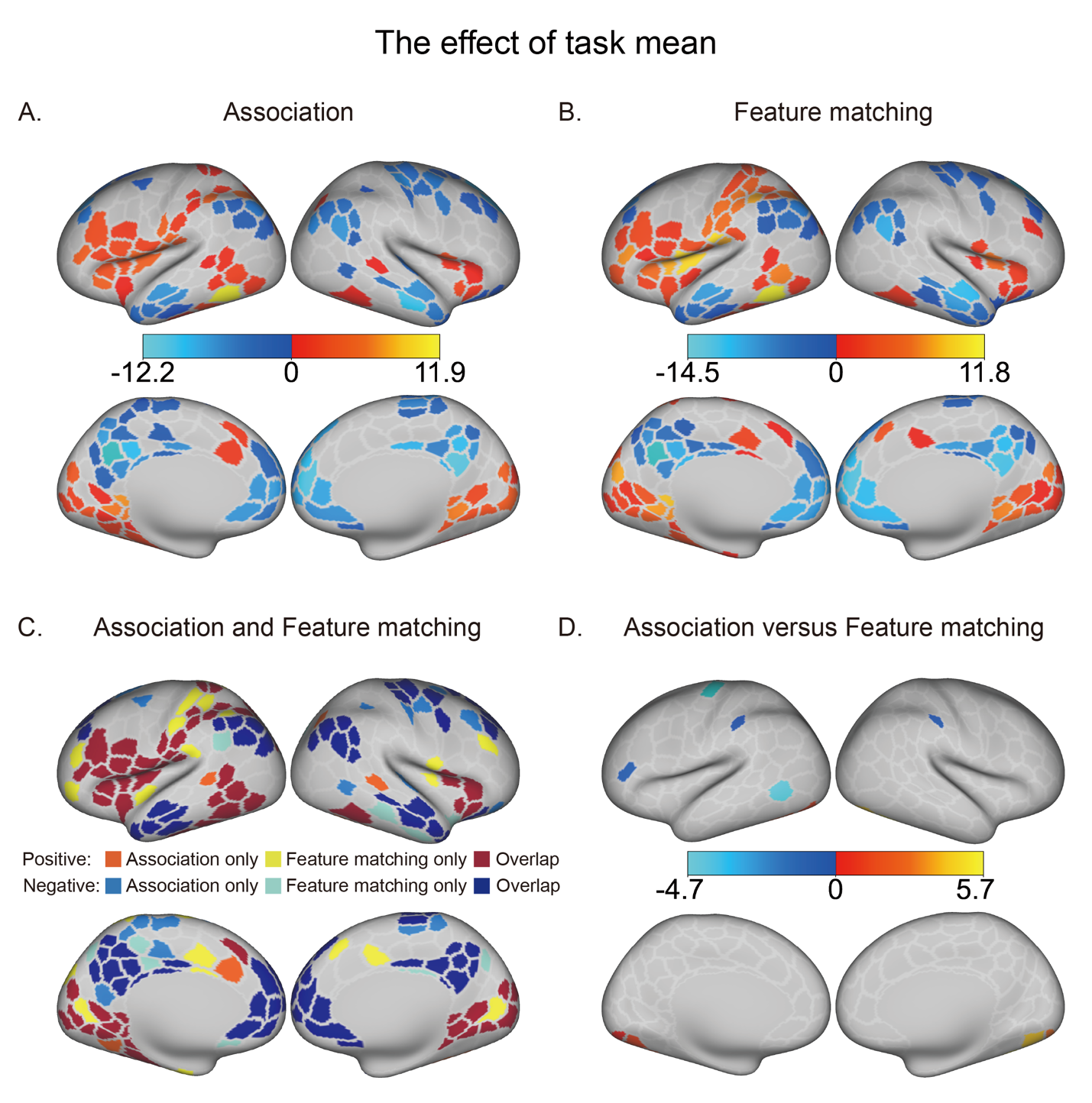

###### Fig. S1. Semantic association and feature matching judgments showed similar patterns of task activation and deactivation. A – The regions showing activation (warm colors) and deactivation (cold colors) in the semantic association task. B – The regions showing activation (warm colors) and deactivation (cold colors) in the feature matching task. C –Overlapping activation and deactivation for the semantic association and feature matching tasks. D – Regions that showed greater activation in the association task compared with feature matching task (warm colors) and the reverse pattern (cold colors).

##### 1.2. The related and non-related trials in the semantic association task showed similar difficulty effects

In the semantic association task, we investigated whether the difficulty effects in related trials were comparable to those in non-related trials by examining how neural responses varied with association strength. Specifically, we analyzed the negative modulation of neural responses by association strength in related trials to identify brain parcels exhibiting greater activation when association strength was weaker for trials judged to be associated. Conversely, for non-related trials, we investigated the positive modulation of neural responses by association strength, aiming to pinpoint parcels showing increased activation when association strength was stronger for trials judged to be non-associated. Fig S2A and S2B depict the difficulty effects of semantic association strength for related and non-related trials, respectively. These effects were significantly correlated (left hemisphere: r = 0.78, p = 0; right hemisphere: r = 0.73, p = 0; spin permutation corrected), confirming their similar difficulty effects.

##### 1.3. The matching and non-matching trials in the semantic feature matching task showed similar difficulty effects

In the semantic feature matching task, we investigated whether the difficulty effects observed in matching trials were analogous to those in non-matching trials, focusing on how neural responses varied with feature similarity. Specifically, we analyzed the negative modulation of neural responses by feature similarity in matching trials. This analysis was aimed at identifying brain parcels that exhibited greater activation when feature similarity was lower, suggesting increased difficulty in these matching trials. Conversely, we assessed the positive modulation of neural responses by feature similarity in non-matching trials. This was to pinpoint parcels that demonstrated heightened activation as feature similarity increased, indicating greater difficulty in non-matching trials. Fig S2C and S2D present the difficulty effects of feature similarity for matching and non-matching trials, respectively. Notably, these effects were significantly correlated (left hemisphere: r = 0.74, p = 0; right hemisphere: r = 0.68, p = 0; spin permutation corrected). This correlation confirms that both trial types exhibited parallel difficulty effects.

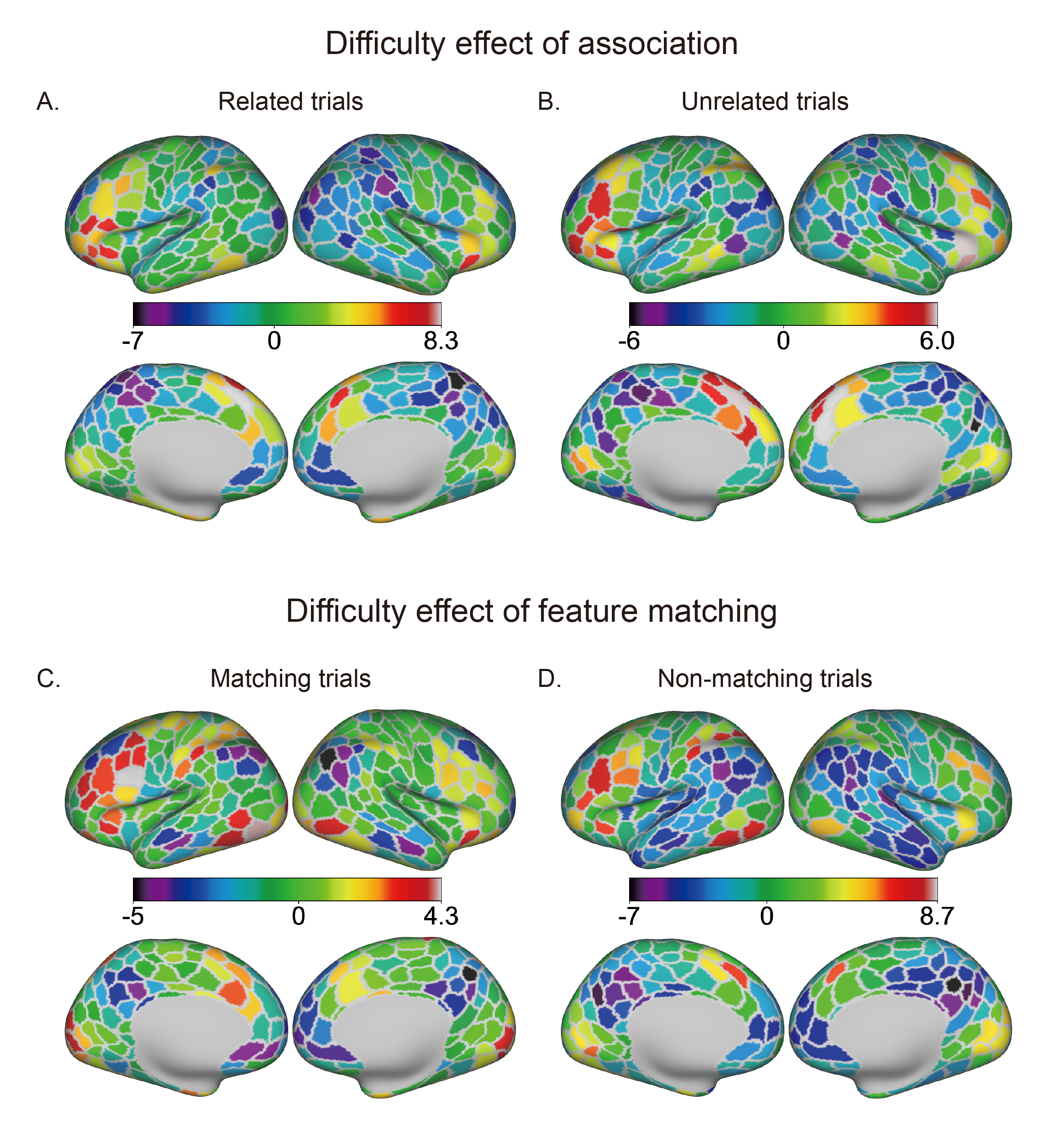

###### Fig. S2. The difficulty effects for related and unrelated trials in the semantic association task and matching and non-matching trials in the semantic feature matching task. A – How neural responses varied with association strength for related trials in the semantic association task. Warm colors mark the brain parcels that exhibited greater activation when association strength was weaker for related trials. B – How neural responses varied with association strength for unrelated trials in the semantic association task. Warm colors highlight the brain parcels showing increased activation when association strength was stronger for unrelated trials. C – How neural responses varied with feature similarity for matching trials in the semantic feature matching task. Warm colors depict the brain parcels that exhibited greater activation when feature similarity was lower for matching trials. D – How neural responses varied with feature similarity for non-matching trials in the semantic feature matching task. Warm colors represent the brain parcels showing increased activation when feature similarity was greater for non-matching trials.
